# Insights into the genetic architecture of resistance to viral haemorrhagic septicaemia virus in rainbow trout from a genome-wide association study to *in vitro* CRISPR-Cas9 functional evaluation

**DOI:** 10.64898/2026.06.09.731144

**Authors:** Valentin Thomas, Bertrand Collet, Edwige Quillet, Michael Marchand, François Huetz, Pierre Boudinot, Florence Phocas, Delphine Lallias

## Abstract

Viral haemorrhagic septicaemia (VHS) is a severe disease affecting rainbow trout (*Oncorhynchus mykiss*) and a wide range of wild freshwater and marine fish species. VHSV threatens rainbow trout aquaculture, as it may cause 100% mortality in fry. Previous studies identified a quantitative trait locus (QTL) on chromosome 3 associated with resistance to VHSV waterborne challenge and reduced viral replication in fin explants, although these findings were obtained using limited genetic diversity.

The objective of this study was to validate and extend the identification of genomic regions associated with resistance to VHSV in the genetically diverse rainbow trout line designated “synthetic.” A genome-wide association study (GWAS) was conducted using whole-genome sequences from parents of progeny classified as resistant or susceptible to a VHSV waterborne challenge. While the QTL on chromosome 3 was not validated in the synthetic line, four novel suggestive SNPs associated with survival following VHSV waterborne challenge were identified on chromosomes 6, 8, 17, and 32. Notably, one SNP on chromosome 17 was located within a gene potentially involved in antiviral defence, a paralog of *lrp1* (low-density lipoprotein receptor-related protein 1).

To further investigate its role, *lrp1* function was analysed *in vitro* using CRISPR-Cas9 genome editing. Three independent *lrp1^-/-^* CHSE-EC cell lines were generated and challenged with VHSV. The results showed that *lrp1* is not essential for viral entry but may modulate the inflammatory response during VHSV infection in epithelial cell lines.

## Introduction

Viral haemorrhagic septicaemia (VHS) is a severe viral disease affecting rainbow trout (*Oncorhynchus mykiss*) and a wide range of freshwater and marine fish species, whether in wild and farmed populations (1), and is listed as a notifiable disease by the World Organisation for Animal Health (WOAH) (1). In rainbow trout, outbreaks of VHS can cause mortality rates approaching 100% in fry, and fish of all sizes are susceptible, with overall mortalities ranging from 5% to 90% (2). The clinical signs of the disease include lethargy, skin darkening, exophthalmia, anaemia, haemorrhages and abdominal swelling caused by oedema. Fish with chronic infections may show no symptoms, and a neurological form of VHS can cause abnormal swimming behaviour, such as flashing and spiralling.

VHS is caused by the viral haemorrhagic septicaemia virus (VHSV), a *Novirhabdovirus* belonging to the *Rhabdoviridae* family (3). The virion is bullet-shaped and enveloped, with a negative-sense, single-stranded RNA genome of approximately 11,000 nucleotides that encodes six proteins: nucleoprotein (N), phosphoprotein (P), matrix protein (M), glycoprotein (G), non-virion protein (NV), and RNA-dependent RNA polymerase (L). The base of the fins represents a major portal for viral entry (4). During the septic phase of infection, the virus spreads widely as it primarily replicates in epithelial and fibroblastic cells, leading to a systemic infection. It is particularly abundant in the kidneys, heart, and spleen. In the chronic stages of the infection, the virus can reach high levels in the brain (5).

Given the absence of therapeutic options, breeding for disease resistance represents a promising approach for VHS control. In this context, the heritability of survival to VHSV in rainbow trout has been estimated in two independent studies at 0.57 (6) and 0.63 (7) on the underlying continuous liability scale when resistance was assessed as a binary trait (dead or alive at the end of a controlled challenge test). A recent study on olive flounder (*Paralichthys olivaceus*) reports a lower heritability of survival to VHSV of 0.145 on the observed binary scale (8).

Moreover, striking differences in susceptibility among rainbow trout isogenic lines further underscore the strong genetic component of resistance (9). QTL mapping studies based on two F2 rainbow trout families, combining the genomes of susceptible and resistant gynogenetic doubled haploid F0 breeders, have identified a major QTL on chromosome 3 associated with both fish survival and viral replication in fin explants (10). This QTL has been confirmed in backcross of highly resistant and highly susceptible isogenic lines (11). One VHS survival-related specific area was also detected in turbot (*Scophthalmus maximus*) LG20, showing a distal syntenic block with the rainbow trout chromosome 3 region where the major QTL was detected (12).

The current study aimed first to confirm, in the genetically diverse rainbow trout line named “synthetic”, the genomic regions previously detected as associated with resistance to VHSV on a narrow genetic basis. The INRAE “synthetic” line was created in the early 1980s by crossing various domesticated populations from France, Denmark, and the United States. This experimental line has since been maintained through full factorial mating, which preserves genetic diversity without intentional selection (13). The QTL detection for the synthetic line was done through a GWAS, using whole genome sequenced parents of progeny either resistant or susceptible to VHSV. Secondly, the role of a potentially relevant candidate gene, *lrp1*, was also tested using CRISPR-Cas9 genome editing in a salmonid cellular model.

## Materials and methods

### Ethic statements

Animal experiments and sampling were performed at the INRAE-IERP fish facilities (doi.org/10.15454/1.5572427140471238E12) in Jouy-en-Josas (France), in accordance with the European Directive 2010/2063/UE. All animal work was approved by the Direction of the Veterinary Services of Versailles, France (building agreement number B78-720) and by the ethics committee of the INRAE Center in Jouy-en-Josas (COMETHEA n°45), France (authorization number #12/053).

### Experimental material

The synthetic challenged fish were obtained by mating 32 females and 60 males from the INRAE synthetic line using a partial factorial design (two full factorials of 11♀x20♂, and one factorial of 10♀x20♂).

VHSV strain 07-71 (serotype 1) isolated from diseased fish from a French rainbow trout farm was propagated in EPC cells as described by Dorson (9). After infection, the cells were incubated at 14 °C in Eagle’s medium supplemented with 2% foetal bovine serum (FBS) and antibiotics (penicillin 100 IU.ml^-1^, streptomycin 0.1 mg.ml^-1^ and kanamycin 0.1 mg.ml^-1^). The virus was harvested when the cytopathic effect was complete.

### Viral infection

A waterborne challenge with VHSV was conducted in 2013 on offspring obtained from mating 32 females and 60 males from the INRAE synthetic line. For the challenge, a total of 2500 offspring (1g) were transferred to a single 300 L tank with a water turnover of 1 volume per hour (10°C) 1 day before infection. Waterborne infections were performed by incubating 1250 fish per tank in two 12 L tanks for 2 h with the virus at 50,000 PFU.ml^-1^ (19/03/2013). Then they were transferred to the previous 300L tank. The challenge lasted 27 days, and mortalities were recorded twice a day. A peak of mortality was observed between 3 and 6 days post-infection, during which 60% of challenged fish died. Furthermore, 12.5% of the fish survived the challenge.

### Genotyping

Fin tissue samples from all dead fish and from survivors at the end of the challenge were preserved in alcohol at -20°C. In total, 552 offspring—comprising 276 first-dead fish and 276 survivors—and 92 parent fin samples were sent to Labogena (https://www.labogena.com/fr/) for genotyping using a panel of 13 microsatellites specific to rainbow trout (S1 Table). Parentage assignment was performed using Vitassign (14), and 96.7 % of offspring were assigned to a unique couple without mismatches.

### Choice of sequenced parents

In a first step, the 49 parents with at least 11 offspring, phenotyped and well-assigned, were ranked by progeny survival rate. Among those parents, 15 were labelled as resistant parents when the progeny survival rate was greater than 80%, and 9 parents as susceptible when the progeny survival rate was equal to or below 20%. On average, 15 progeny were phenotyped and well assigned to each of the 49 parents. In a second step, 7 parents presenting an over-representation of crosses with resistant or susceptible mates were discarded for further analysis.

Among the remaining 42 parents, a total of 14 individuals were selected for whole genome sequencing as the parents with the more extreme phenotypes based on their progeny survival rates: 7 resistant parents (with 83-100% offspring survival) and 7 susceptible parents (with 0-20% offspring survival) (Fig 1).

**Fig 1.**
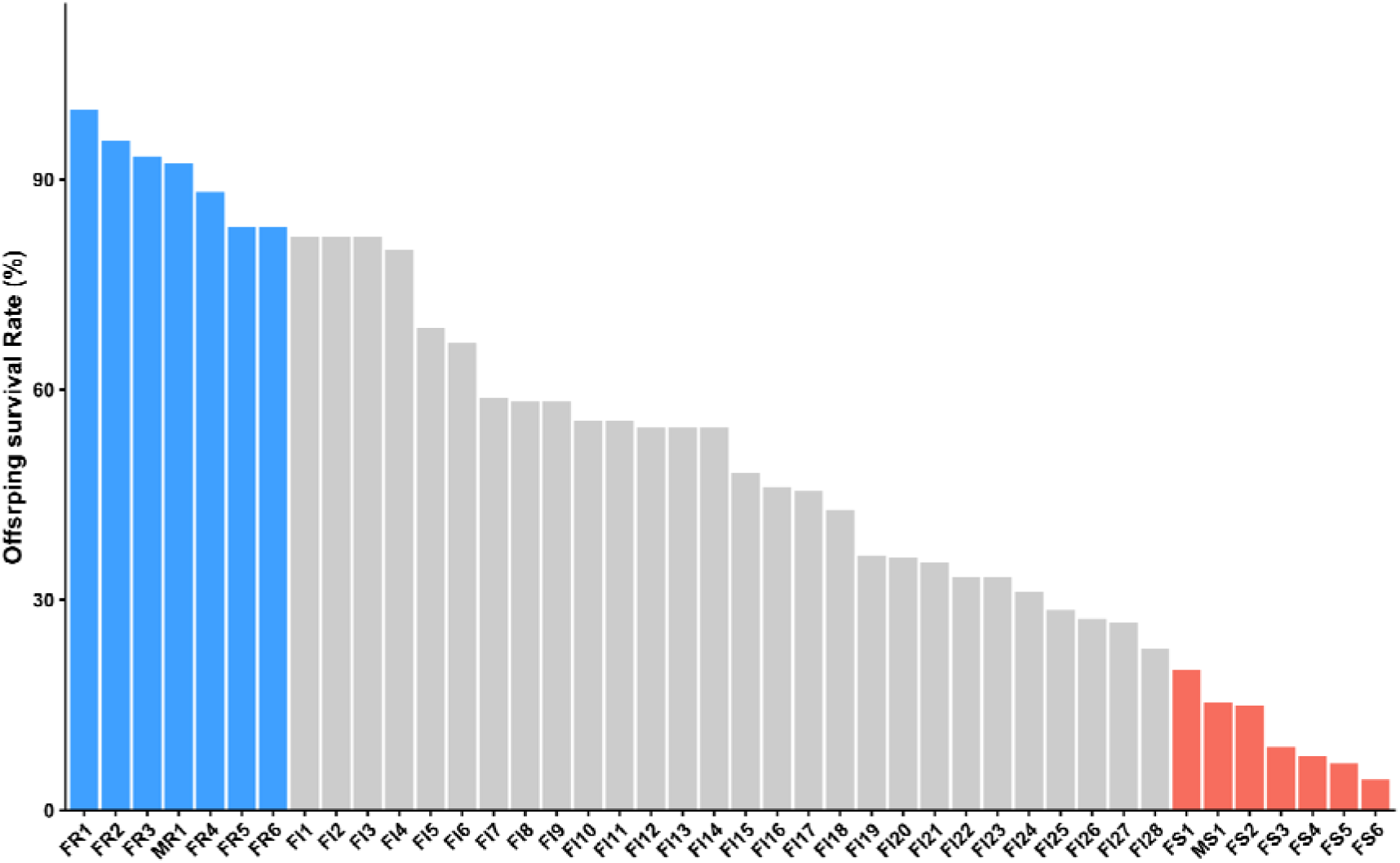
Strategy for selecting extreme parents for whole genome sequencing, based on offspring survival rate. Distribution of offspring survival rates (%) across the 42 parents. Coloured bars indicate the selected extreme groups: parents with offspring survival rates ≥ 82.6% (blue) are classified as resistant, while those with survival rates ≤ 20% (red) are classified as susceptible.

### Sequencing

The 14 fish fin samples (20 mg) were ground with Precellys® Evolution Touch (6 beads, successively 6000 rpm 2×20sec, 6000 rpm 2×40sec and 10 000 rpm 2×40sec). DNA extraction was carried out using Wizard Genomic DNA Purification Kit from Promega. DNA libraries from all individuals were prepared using the TruSeq Nano Kit. Sequencing in paired-end 2×150 bp was performed in a single lane on a high-throughput Illumina NovaSeq 6000 sequencer at the INRAE sequencing platform Get-Plage in Toulouse France (https://doi.org/10.15454/1.5572370921303193E12), and reached an average of 22.4X.

### Variant calling

Variant calling was done with nf-core/sarek v3.4.0 (15,16) with a bootstrap step. Sequences were aligned to the rainbow trout reference genome USDA_OmykA_1.1, Assembly GCA_013265735.3. Called SNP during bootstrap underwent quality control filtering based on GATK best practices. The criteria for filtering included: FS ≥ 60.0, QD ≤ 2.0, MQ ≤ 40.0, SOR ≥ 3.0, MQRankSum ≤ -12.5, ReadPosRankSum ≤ -8.0, and Qual < 30.0. Additionally, non-nuclear, non-variant and multi-allelic SNP with missing genotypes were excluded, resulting in a bootstrap panel of 18,672,695 SNP.

To distinguish true variants from sequencing artefacts, the SNP bootstrap panel was used for the recalibration step. The recalibrated BAM files were then used as input for the second variant calling step. The SNP identified during this second step were further filtered according to the same GATK guidelines. Therefore, SNP that were non-nuclear, non-variant, multi-allelic, had a minor allele frequency below 10.7% (less than 3 minor alleles over 28), or had more than 10 missing genotypes were excluded from the analysis. Ultimately, a list containing 13,219,207 SNP was used for further analysis.

### GWAS

A QTL detection was performed using the genotypes of 13,219,207 SNP for the 14 parents of the synthetic line with extreme progeny survival rates.

Pairwise relationships among all individuals were assessed by calculating probabilities of Identity by State using the plink1.9 --genome parameter (17). Most individuals were not or weakly related (P_IBS_ ≤ 0.111); however, 2 pairs —MS1/FS2 and FS1/FS5— share approximately 30% of their alleles, suggesting they may be half-siblings (S2 Table). To avoid any family structure that could bias GWAS results, the analysis was conducted across 4 subsets of 12 low-related fish among the 14 sequenced individuals (P_IBS_ ≤ 0.111). From each pair of related fish, a single individual was included in a subset, with all possible combinations distributed across 4 subsets numbered 1, 2, 3, and 4.

This procedure resulted in 4 analyses of 12 low-related individuals, each comparing 7 resistant to 5 susceptible fish. Hence, GWAS were performed SNP by SNP using Fisher’s exact test (R). To get significance thresholds, the “simple*M”* method was applied to account for multiple tests and correlations between close SNPs, applied iteratively in 10kb windows across the genome as described in Gao et al. (2008). The effective number of independent tests in each window was calculated as N_eff_ = number of SNP explaining 99% of the variation in the window. Accordingly, the effective number of tests along the genome was N_eff_ = 966,004. The genome-wide significance threshold corresponding to a Bonferroni-like correction for a p-value of 0.05 was then derived as -log_10_(0.05/N_eff_) = 7.3. A second threshold was defined to indicate SNP with a suggestive effect when -log_10_(0.05/N_chr_) ≥ 5.81. This threshold corresponds to the Bonferroni-like correction taking the median effective number of tests derived per chromosome (N_chr_ = 32,345).

QTL were considered as ‘true’ as soon as they were detected in 2 subsets out of the 4 analyses. SNP were said to be significant if at least 2 tests gave values above 7.3, and suggestive if at least 2 tests gave values above 5.81.

For all significant or suggestive SNP, linkage disequilibrium (LD) was calculated with plink1.9 --r2 inter-chr and --ld-window-r2 0 parameters (Chang et al., 2015), between these top SNP and all other SNP within a region of 1Mb (± 500kb around a top SNP). All of these SNP were annotated using VEP version 114 (McLaren et al., 2016). To further improve the annotation process, genes were compared against the human genome, retaining the symbol of the gene that showed the best BLAST hit as the annotation (19). To identify these optimal BLAST hits, the rainbow trout proteome was obtained and filtered to include only the longest isoform for each gene. Protein sequences underwent a BLASTP analysis using Diamond v2.0.9.147 (20) against the human (GCA_000001405.28) and zebrafish (GCA_000002035.4) peptide sequences sourced from Ensembl (21). All parameters were set to their default values.

### Cell culture condition

The Chinook salmon *(Oncorhynchus tshawytscha*) embryo (CHSE-EC) cell line (Cellosaurus CVCL_DG46) and all derivatives were grown in Leibovitz’s L-15 medium (Gibco) supplemented with 10% FBS (Eurobio) and penicillin (100 U.mL^-1^)-streptomycin (100 μg.mL^-1^) (BioValley), 500 μg.mL^-1^ G418 (Invivogen), 30 μg.mL^-1^ hygromycin B Gold (Invivogen). All cell lines were maintained at 20 °C without CO_2_.

### Development and validation of *lrp1-/-* cell lines

The CHSE-EC cell line was used to develop a *lrp1^-/-^*cell line. According to the Ensembl 115 release, the gene *lrp1*, detected in the GWAS, has 3 paralogs in rainbow trout and 4 in chinook salmon. They are all co-orthologs of the human *LRP1* (ENSG00000123384).

Two single guide RNAs (sgRNAs) were designed to target all four paralogs of *lrp1* in chinook salmon using the CRISPOR v5.2 web tool (22). Both sgRNA specifically target the following genes: ENSOTSG00005004145 (exon 8), ENSOTSG00005039789 (exon 9), ENSOTSG00005041107 (exon 8), and ENSOTSG00005058412 (exon 9), without any mismatches. The cutting sites for the two sgRNAs are located 13 base pairs apart. No off-target sites with fewer than three mismatches in the first 12 bp adjacent to the protospacer adjacent motif (PAM) were found in the chinook salmon genome (*Oncorhynchus tshawytscha* NCBI GCA_018296145.1 Otsh_v2.0), thereby confirming the specificity of the sgRNAs. Additionally, a sgRNA targeting the mEGFP gene was used for screening (sens TCCTAATACGACTCACTATAGGCGAGGGCGATGCCACCTAGTTTTAGAGCTAGAAATAGCAAGTTAAAATAAGGCTAGTCCGTTATCAACTTGAAAAAGTGGCACCGAGTCGGTGCTTTT, antisens AAAAGCACCGACTCGGTGCCACTTTTTCAAGTTGATAACGGACTAGCCTTATTTT AACTTGCTATTTCTAGCTCTAAAACTAGGTGGCATCGCCCTCGCCTATAGTGAGTCGTATTAGGA).

The sgRNAs were synthesized using the T7 RiboMAX Express Large Scale RNA Production System kit (Promega) according to the manufacturers’ instructions using 3.8 μg of each sens (1-TCCTAATACGACTCACTATAGTTCCCCCACGGTATCACCCGTTTTAGAGCTAGAAATAGCAAGTTAAAATAAGGCTAGTCCGTTATCAACTTGAAAAAGTGGCACCGAGTCGGTGCTTTT/ 2- TCCTAATACGACTCACTATAAGGTCCAGGGTGATACCGTGGTTTTAGAGCTAGAAATAGCAAGTTAAAATAAGGCTAGTCCGTTATCAACTTGAAAAAGTGGCACCGAGTCGGTGCTTTT) and antisens (1-AAAAGCACCGACTCGGTGCCACTTTTTCAAGTTGATAACGGACTAGCCTTATTTTAACTTGCTATTTCTAGCTCTAAAACGGGTGATACCGTGGGGGAACTATAGTGAGTCGTATTAGG/ 2- AAAAGCACCGACTCGGTGCCACTTTTTCAAGTTGATAACGGACTAGCCTTATTTTAACTTGCTATTTCTAGCTCTAAAACCACGGTATCACCCTGGACCTTATAGTGAGTCGTATTAGGAA) RNA that spontaneously annealed as the dsDNA template. The RNA synthesis mix was incubated 1h at 37°C and then with 1 μL of RQ1 DNase (Promega) for 30 h at 37 °C and purified using TRIzol^TM^ reagent (Invitrogen), according to the manufacturers’ instructions. The sgRNAs were resuspended in RNase- and DNase-free water and quantified using a Nanodrop spectrophotometer. The ability of each sgRNA to cut the target sequence was confirmed by *in vitro* efficiency assay. Genomic DNA was extracted from T25 confluent CHSE-EC cells using NucleoSpin Tissue Mini kit (Macherey–Nagel), according to the manufacturer’s instructions. Genomic DNA segments containing the targeted sites were amplified by PCR using Q5 Polymerase (Promega) with the following primers (Table 1).

**Table 1.**
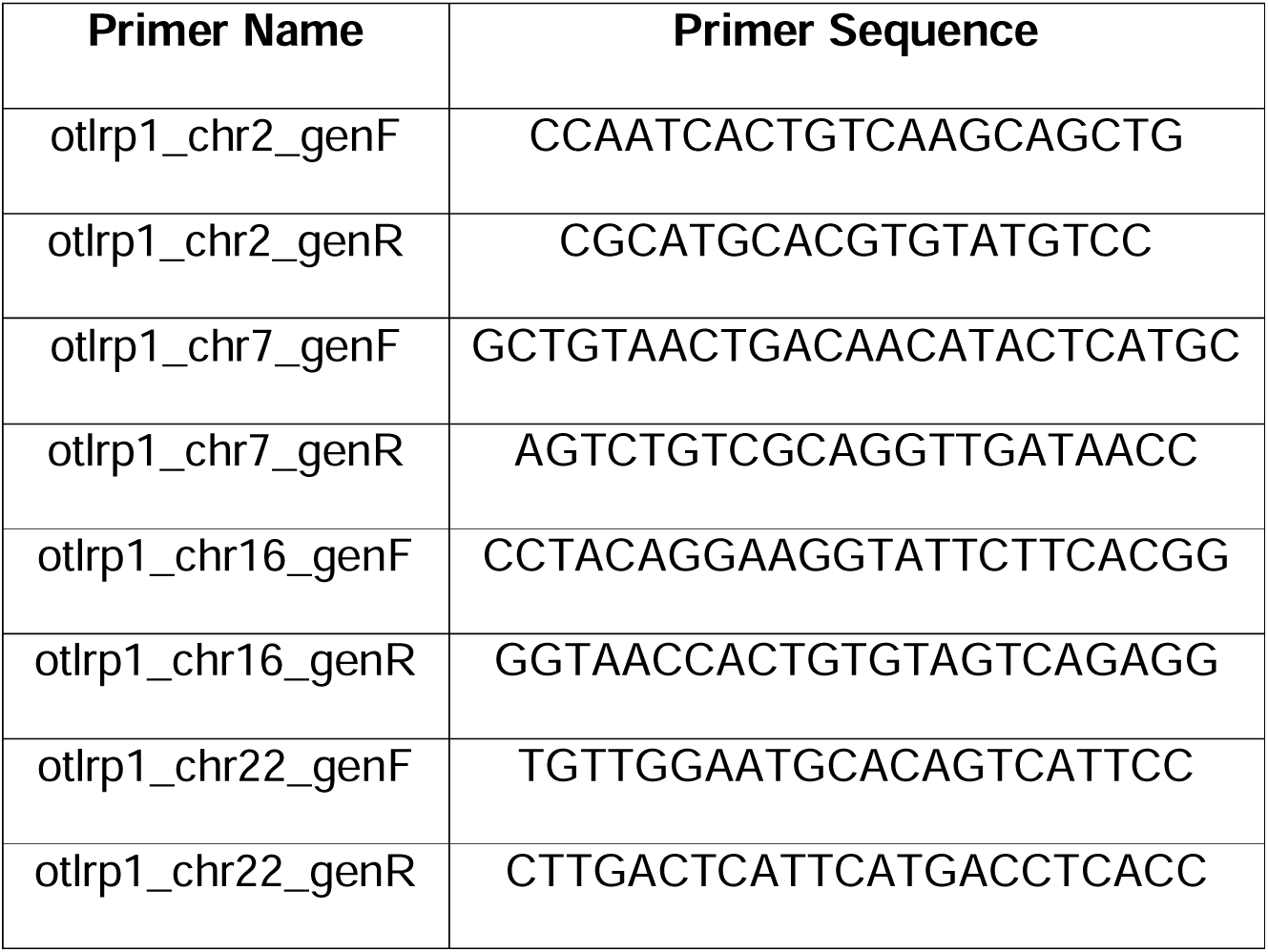
List of *lrp1* paralogs specific primers in chinook salmon.

The PCR cycling program was performed in a thermal cycler (Eppendorf) and was as follows: 98 °C for 30 s then 35 cycles of 98 °C for 10 s, 60 °C for 30 s, 72 °C for 1min, and a final extension of 72 °C for 2 min. The PCR products were purified using NucleoSpin Gel and PCR Clean-up Mini kit (Macherey–Nagel). After electrophoresis validation of length and purity (200mL TAE 1X, 2% agarose, 4μl Gel Green® 10,000X (Merck), 1h 100V migration), PCR products were sequenced (Sanger sequencing service, Eurofins). After sequence analysis and evidence for genome editing at the cell population level, the cells were passaged a second time, and a small proportion was seeded at very low density (10-fold serial dilutions) on a 6-well plate. Clonal cell patches were identified and examined under a fluorescent microscope (Zeiss Axio Observer Z1, Oberkochen, Germany fluorescent inverted microscope). mEGFP-deficient (non-fluorescent) clones were selected, detached mechanically by gently scraping with pipette tip while aspirating at the same time and sub-cultured in separate 25cm^2^ flasks (15 clones). Of the clones isolated with this method and propagated, one was validated for *lrp1* mutation as described above. To enrich the clones catalogue cells were also FACS-sorted. Cells were detached by trypsin-EDTA action, and mEGFP-deficient single cells were individualized by a BD FACS Aria Fusion flow cytometer (Institut Pasteur, Paris, France) using a 100-mm nozzle into a 96-well plate (Sarstedt) in L-15 medium (Gibco) supplemented with 10% FBS (Eurobio) and penicillin (100 U.mL^-1^)-streptomycin (100 μg.mL^-1^) (BioValley), 500 μg.mL^-1^ G418 (Invivogen), 30 μg.mL^-1^ hygromycin B Gold (Invivogen). Two months later, surviving clones were subcultured and propagated, from which two clones were isolated and validated for *lrp1* mutation as described above.

### Tissue culture infectious dose (TCID_50_) assay

To clarify the potential impact of *lrp1* deletion on survival, a TCID_50_ assay was conducted. CHSE-EC (WT) and CHSE-EC-*lrp1*^⁻/⁻^ clonal lines were seeded in 96-well plates in quadruplicate at a final density of 45,000 cells per well in L-15 medium supplemented with 10% FBS and incubated at 20 °C until the cells reached at least 80% confluency. Cells were then infected with VHSV 07.71 at 910 PFU.mL^-1^ in the first column, followed by 4-fold serial dilutions across the plate. The last column was left uninfected and served as a control. Plates were subsequently incubated at 14 °C for 7 days. Cells were fixed with 4% paraformaldehyde for 15 min, washed, and allowed to dry for 1 hour at room temperature. The monolayer was then stained with 50 µL of 0.4% crystal violet for 15 min, washed, and left to dry overnight. TCID_50_ was estimated by scoring wells according to cytopathic effect (CPE): wells with ≤ 50% of the cell monolayer showing CPE were considered negative (0), whereas wells with > 50% CPE were considered positive (1). For each cell line, infection outcome (0/1) was modelled as a function of log_10_-transformed viral dilution using logistic regression (binomial GLM with logit link). The model was fitted using all individual measurements across three biological replicates per cell line with the glm() function in R. For each cell line, TCID_50_ was calculated as the concentration yielding 50% infection probability (ie -intercept / slope), with 95% confidence intervals derived using the delta method. Statistical comparisons between clones and WT cells were performed using likelihood ratio tests (LRT) on nested models, testing whether allowing separate dose-response parameters significantly improved model fit. P-values were adjusted for multiple testing using the Benjamini-Hochberg (BH) correction, with significance threshold set at adjusted p < 0.05. All analyses were conducted in R.

### qPCR

In order to test the effect of *lrp1* deletion on different viral responses, the expression of key inflammatory genes as well as viral RNA expression were measured in WT and *lrp1^-/-^* cell lines post-infection. CHSE-EC (WT) and CHSE-EC-*lrp1*⁻*/*⁻ clonal lines were seeded in 24 well plates (300,000 cells/well, in triplicate) in L-15 medium supplemented with 10% FBS and incubated at 20 °C to ≥80% confluency. Cells were infected with VHSV 07.71 at a multiplicity of infection of one (MOILJ=LJ1) and incubated at 14 °C for 30 h. After removing the medium, cells were collected in 500LJµL TRIzol and stored at −80 °C. RNA was extracted by adding chloroform (100 µL), centrifuging at 12,000 g for 15 min at 4 °C, and transferring 250 µL of the aqueous phase to fresh tubes. RNA was precipitated with 250 µL isopropanol supplemented with GlycoBlue, washed with 70% ethanol, dried, and resuspended in 50LJµL RNase-free water. First-strand cDNA was synthesised from 385 ng RNA using a Takara PrimeScript RT Master Mix cDNA kit. First-strand cDNA samples were diluted fourfold (working stock) with RNase/ DNase-free water and stored at −20 °C. Real-time PCR (qPCR) analyses were performed with a Bio-Rad CFX Duet Mastercycler and BIORAD - CFX Connect. All assays were performed in 25 μL reactions in 96-well plates, in duplicate. Each reaction mix contained 2 μL of cDNA, 12.5 μL of Takara TB Green Premix Ex Taq, and 10.5 μL of a forward and reverse primer solution, achieving a final concentration of 200 nM for each primer. PCR cycling conditions were 1 cycle of 95 °C for 1 min, followed by 40 cycles of 95 °C for 10 s and then 60 °C for 30 s. Melting curve analysis (thermal gradient from 60 to 95 °C) was then used to confirm the amplification of a single product. Each plate also included “no template” negative controls in quadruplicate (cDNA replaced with water). Efficiency was calculated for each primer from a serial dilution PCR. *vhsv g* expression was used to compare levels of viral propagation across cellular mutant clones. The upregulation of inflammatory genes was assessed by *ikb* and *il1b* expression. All target gene expression was normalised to *rps29*. Relative expression levels were calculated for each cell line, comparing infected and non-infected wells. All primers used in this study are available in S3 Table.

## Results

### Selection of resistant and susceptible fish

To detect genomic regions associated with resistance to VHSV, the 14 most extreme parents were selected for whole-genome sequencing based on their offspring’s mean survival rate in a waterborne challenge to VHSV. The average offspring survival rate of the 42 candidate parents was 50.5%. This selection process categorised the extreme parents into two groups: a resistant group (7 fish), with a mean offspring survival rate of 91% and 17 progeny well-assigned on average, and a susceptible group (7 fish), with a mean offspring survival rate of 11% and 16 progeny well-assigned on average (Fig 1).

### Detection of four new suggestive SNP correlated with VHSV resistance

No SNP was found to be significantly correlated with VHSV resistance in any of the four subsets of low-related parents. However, five top SNP reached the suggestive threshold of 5.81 in at least two subsets. They are located on chromosomes 6, 8, 17, and 32, as illustrated in Fig 2.

**Fig 2.**
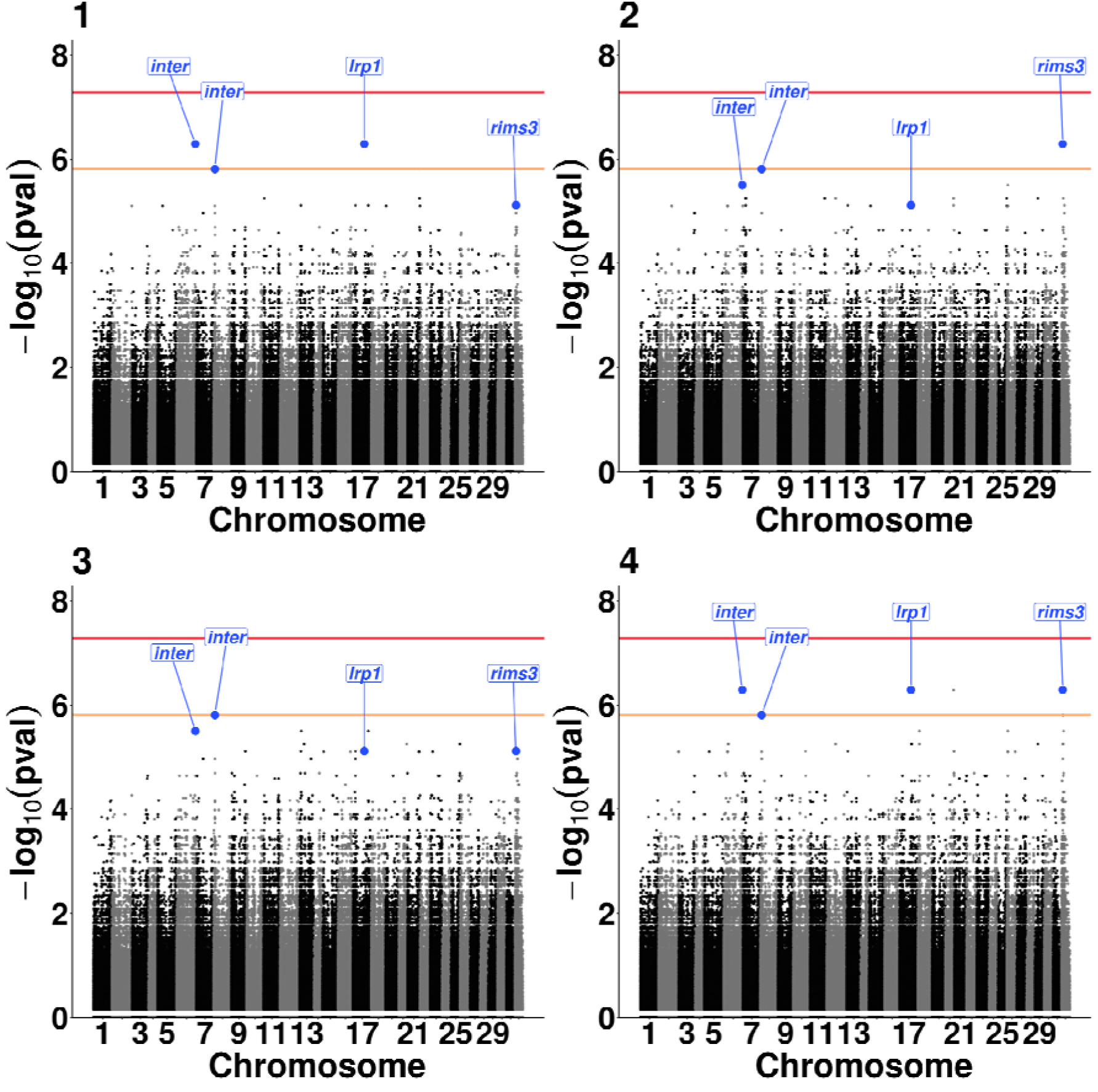
Results of the GWAS conducted across four subsets of synthetic extreme parents. The red line indicates the genome-wide significant threshold (7.29), while the orange line represents the chromosomal suggestive threshold (5.81). Top SNP are labelled by blue dots (−log_10_(pval) >= 5.81 in at least two subsets). The different chromosomes are represented alternatively in black and grey.

Among the suggestive SNP, three are intergenic, two of which are located only 4 base pairs apart on chromosome 8. One suggestive SNP is located within an intron of the *lrp1* gene (ENSOMYG00000036165), which encodes the *low-density lipoprotein receptor-related protein 1* (CD91). This protein is a multifunctional cell-surface receptor belonging to the LDL receptor family. Another suggestive SNP is found within an intron of the *rims3* gene (ENSOMYG00000050898), which encodes *regulating synaptic membrane exocytosis 3*, involved in the binding of transmembrane transporters at the presynaptic active zone (23). All information and precise location of suggestive SNP are shown in Table 2.

**Table 2.** Summary of top SNP reaching the suggestive significance threshold.

| Chromosome | Position (bp) | MA F | Consequence | Rainbow trout gene | Homo sapiens gene | logp 1 | logp 2 | logp 3 | logp 4 |
| --- | --- | --- | --- | --- | --- | --- | --- | --- | --- |
| 6 | 96,729,959 | 0.46 | Intergenic | - |  | 6.29 | 5.50 | 5.50 | 6.29 |
| 8 | 6,684,051 | 0.46 | Intergenic | - |  | 5.81 | 5.81 | 5.81 | 5.81 |
| 8 | 6,684,055 | 0.46 | Intergenic | - |  | 5.81 | 5.81 | 5.81 | 5.81 |
| 17 | 58,951,960 | 0.46 | Intronic | ENSOMYG00000036165 | LRP1 | 6.29 | 5.12 | 5.12 | 6.29 |
| 32 | 5,974,340 | 0.46 | Intronic | ENSOMYG00000050898 | RIMS3 | 5.12 | 6.29 | 5.12 | 6.29 |

MAF = Minor Allele Frequency, logp n = $\log(p\text{-val})$ in subset n

### Fine mapping of the QTL regions

To characterize the genomic regions associated with the top SNPs and prioritise putative causal variants, the four QTL regions were analysed. For each suggestive SNP, variants in strong linkage disequilibrium (r² ≥ 0.85) within a 1 Mb window (±500 kb around the top SNP) were identified (S4 Table).

Among the SNPs in strong linkage disequilibrium with the suggestive variants, several were located within genes across the corresponding 1 Mb regions (S4 Table). The potential candidate genes identified included *rhpn2, prx, slc7a10, lrp3, wdr88, gpatch1, muc4,* and *ttn* on chromosome 6; *dll4, vps18,* and *zfyve1* on chromosome 8; *lrp1* and *esyt1* on chromosome 17; and *colq, rims3, nhsl3,* and *mfsd2a* on chromosome 32.

### *lrp1* phylogeny and CHSE-EC-lrp1^-/-^ genotypes

Among the top suggestive SNP, only two were located inside genes, either *lrp1* on chromosome 17 or *rims3* on chromosome 32, making them potential candidate genes for *in vitro* functional validation. While there was no evidence for the potential role of *rims3* during viral infection, *lrp1* has been shown to be implicated in host resistance to viral infection in mammals. Indeed, *lrp1* is involved in rhabdovirus and other viruses’ entry mechanisms in mammals (24). Moreover, *lrp1* affects extracellular matrix dynamics, especially fibronectin recycling (25). It is also known that VHSV binds to fibronectin, suggesting *lrp1* may be an entry cofactor in receptor-mediated endocytosis (26).

Consequently, we tested if *lrp1* could contribute to cell survival during VHSV infection, potentially by regulating viral entry.

The CHSE-EC cell line was selected as an *in vitro* model to test this hypothesis. This epithelial-like cell line is derived from Chinook salmon and belongs to the same genus (*Oncorhynchus*) as rainbow trout. Importantly, CHSE-EC cells are susceptible to VHSV infection (27–29). In addition, CHSE-EC was derived from the CHSE-214 cell line as a tool for efficient screening of genome edit clones using CRISPR/Cas9 in fish cells (30). This cellular model was therefore chosen for knock-out experiments, enabling direct assessment of the role of *lrp1* in VHSV resistance.

Four paralogs of *lrp1* are in fact present in the chinook salmon genome. In addition to the gene identified in the GWAS on rainbow trout chromosome 17, which is located on chinook salmon chromosome 2, 3 other paralogs are present on chinook salmon chromosomes 7, 16 and 22 (Fig 3). Synteny analysis indicates that these genes correspond to *lrp1* family members located on rainbow trout chromosomes 17, 9 and 16, respectively. It has also been shown that all lrp1 paralogs are expressed in CHSE-EC, with the chromosome 2 paralog being the most expressed (31).

**Fig 3.**
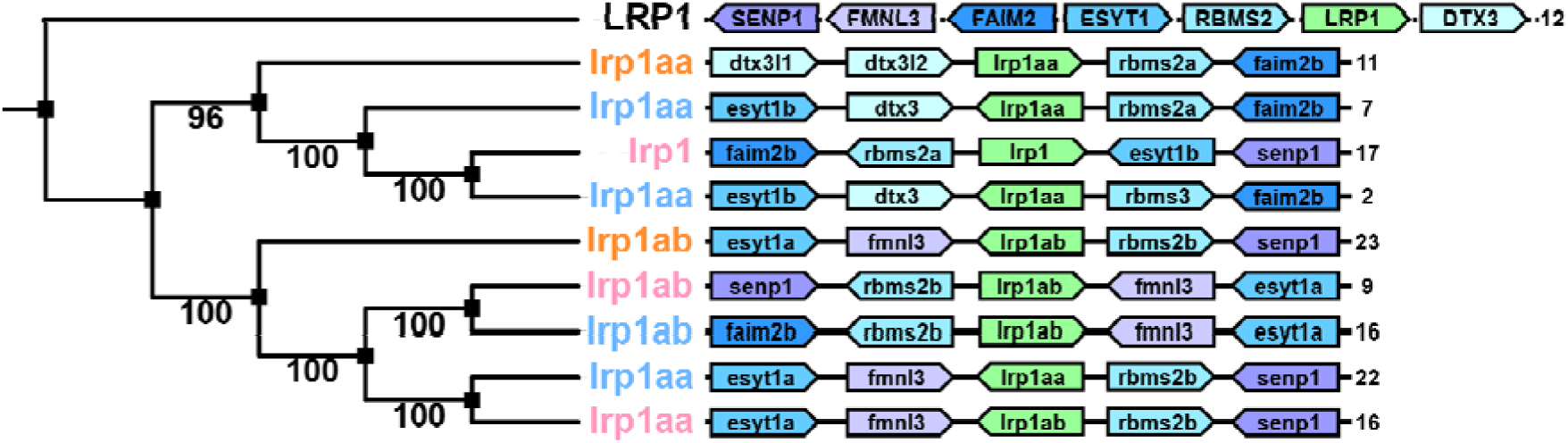
Phylogenetic Maximum Likelihood tree of *lrp1* with 100 bootstraps. The tree with highest log likelihood is shown (−30,717.23). The percentage of replicate trees in which the associated taxa clustered together (100 replicates) is shown next to the branches. The different *lrp1* across species are shown as following: human (black, ENSP00000243077), zebrafish (orange, ENSDARP00000133829 and ENSDARP00000082641 in descending order), chinook salmon (blue, ENSOTSP00005153585, ENSOTSP00005007655, ENSOTSP00005124652 and ENSOTSP00005155688 in descending order) and rainbow trout (pink, ENSOMYP00000079831, ENSOMYP00000028996 and ENSOMYP00000128581 in descending order). Evolutionary analyses were conducted in MEGA12. Chromosome and synteny of each gene are shown on the right.

To test the impact of *lrp1* on VHSV infection, loss of function experiments were performed, targeting chinook *lrp1* family members using a CRISPR-Cas9 approach. Three independent clones (C15, C19, C24) were produced, all edited by the same sgRNA 2. All mutated clones carried premature stop codons in the paralogs on chromosomes 2 (counterpart of rainbow trout *lrp1* detected in the GWAS) and 22, resulting in truncated, nonfunctional proteins (Table 3). Various mutations were found in the remaining paralogs (on the chromosomes 7 and 16) between the three cellular clones generated (Table 3). Sequences of each clonal line at the *lrp1* loci are provided in S5 Table.

**Table 3.**
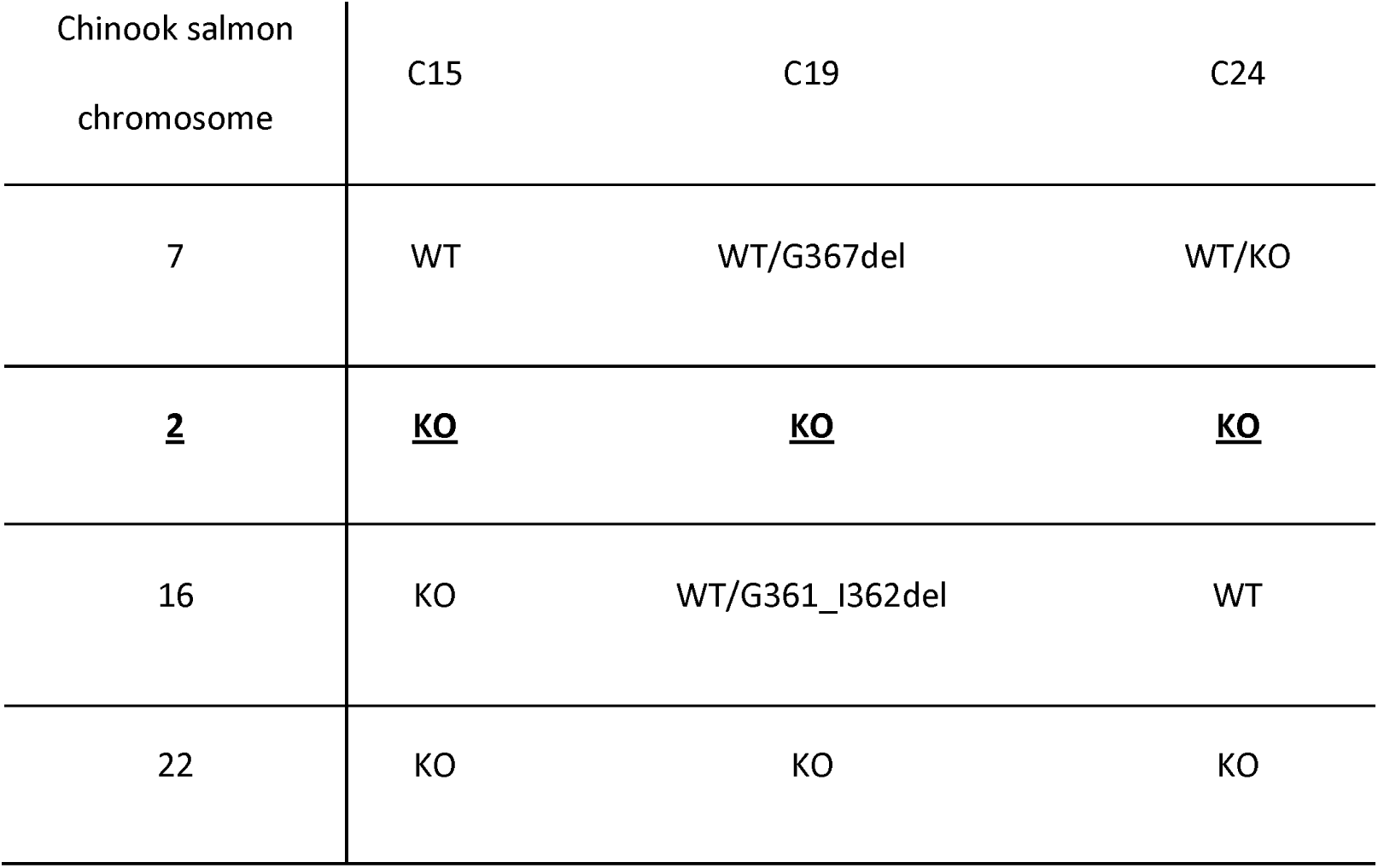
**Three CHSE-EC-*lrp1^−/−^* genotypes for the four paralogs**, with the GWAS *lrp1* chinook ortholog in bold. WT refers to no edition relative to genome and protein modifications at sgRNA 2 CRISPR target site are reported in case of CRISPR edition.

| Chinook salmon<br>chromosome | C15 | C19 | C24 |
| --- | --- | --- | --- |
| 7 | WT | WT/G367del | WT/KO |
| **<u>2</u>** | **<u>KO</u>** | **<u>KO</u>** | **<u>KO</u>** |
| 16 | KO | WT/G361_I362del | WT |
| 22 | KO | KO | KO |

### The deletion of *lrp1* does not confer resistance to VHSV in CHSE-EC

The impact of *lrp1* disruption was first evaluated by VHSV titration in the supernatant of the edited clones after experimental infection. The TCID_50_ assay indicated that the deletion of *lrp1* did not confer protection against VHSV in CHSE-EC. Although mean TCID_50_ (pfu.mL^-1^) differed among WT (22.5), C15 (32), C19 (6.4), and C24 (9), differences between *lrp1^-/-^* lines and WT were not statistically significant (C15 and C24); the *lrp1^-/-^* C19 line even showed lower survival (C19) to VHSV than the WT (Figure 4AB). Also, VHSV-G expression levels 30 h post-infection (MOIL=L1) were comparable between WT and the 3 knockout clones (Fig 4C). Altogether, these data suggested that CHSE-EC survival is not solely dependent on the *lrp1* paralog on chromosome 2 (Fig 4AB). However, disruption of *lrp1* and its paralogs may affect the induction of inflammation by VHSV infection, which is correlated with the regulation of *ikb* and *il1b* expression. Both C19 and C24 showed more pronounced down-regulation of *ikb* than WT, and C19 exhibited a significantly stronger induction of *il1b* (Fig 4C).

**Fig 4.**
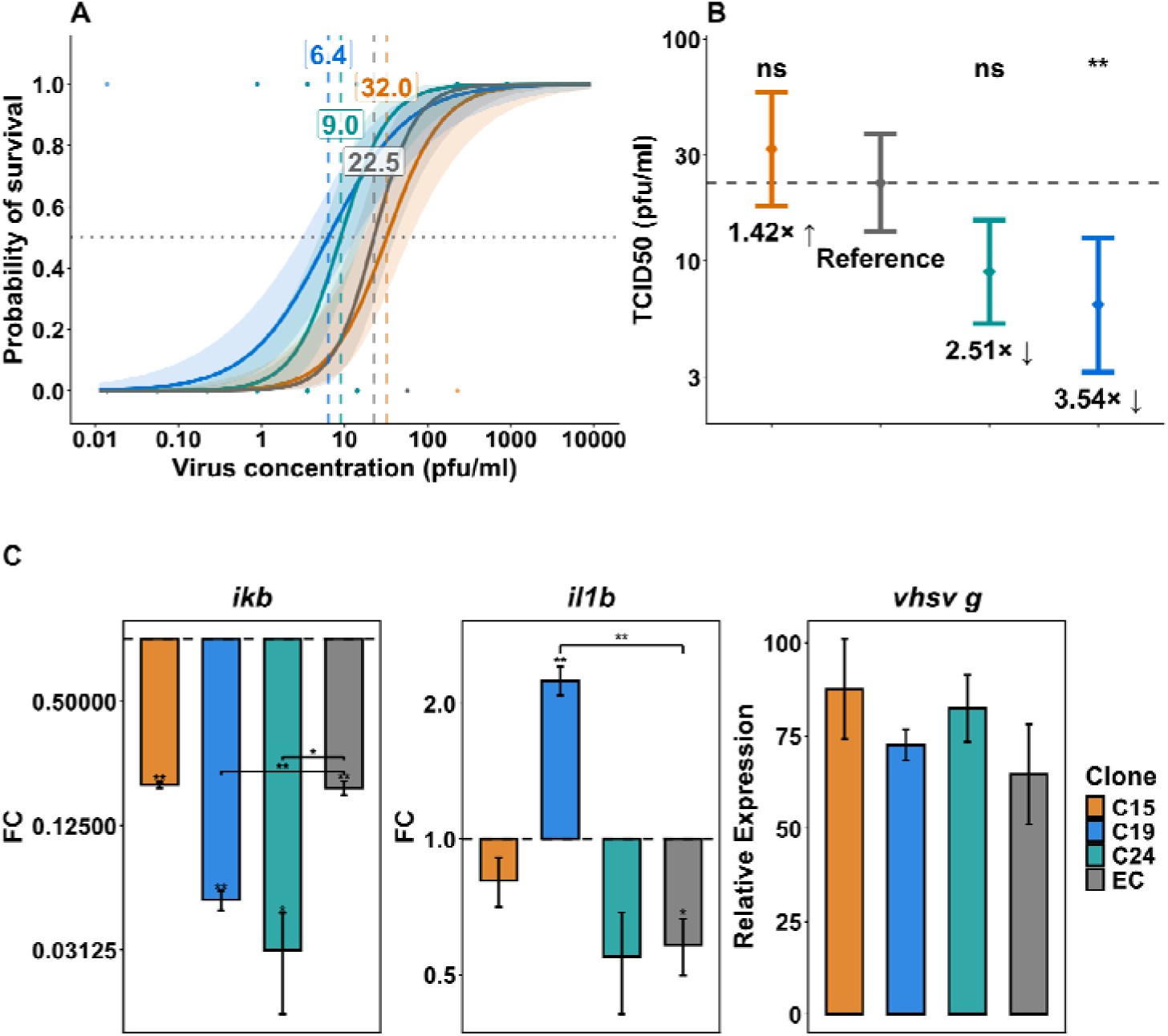
**A. Dose–response curves showing the probability of survival as a function of virus concentration (pfu.mL^-1^) for each cell line (n=3 biological replicates).** Points represent individual experimental measurements from three independent biological replicates per cell line, solid lines represent fitted logistic regression models, using all individual measurements across three biological replicates per cell line, and shaded areas indicate 95% confidence intervals (delta method). Dashed vertical lines indicate estimated TCID_50_ values (50% survival probability). **B. Plot summarising TCID_50_ estimates for each cell line (n=3 biological replicates).** Points represent maximum likelihood estimates from logistic regression, and horizontal bars indicate 95% confidence intervals (delta method). The dashed vertical line indicates the WT reference. Fold-changes relative to WT and statistical significance of differences (Likelihood Ratio Test with Benjamini–Hochberg correction) are indicated next to each comparison. C. Integrated summary of *ikb* and *il1b* expression (left and center panels) and viral gene expression (right panel) measured by qPCR. *ikb* and *il1b* expression is shown as fold change relative to control samples (2^-ΔΔCt^), mean values represented by bars, and error bars indicating standard deviation across 3 technical replicates. Statistical significance versus control was assessed using one-sample t-tests on log_2_FC, with Benjamini–Hochberg correction for multiple testing. Brackets indicate pairwise comparisons between WT and clone conditions when significant. Viral expression (right panel) shows relative quantification of *vhsv g* normalised to the reference gene *rps29*. Bars represent mean relative expression per sample, with error bars indicating standard deviation across 3 technical replicates.

## Discussion

This study aimed to enhance understanding of the genetic basis of VHSV resistance in rainbow trout by combining two approaches: (1) a GWAS utilising whole genome sequencing of extreme parents from the synthetic line, which has a broader genetic background than previous studies, and (2) *in vitro* functional evaluation of *lrp1* family members’ influence on VHSV resistance.

It was assessed whether sequencing a limited number of individuals with extreme phenotypes could provide sufficient power for GWAS at very high SNP density (∼13 million markers). With this method, several suggestive genomic regions were identified on chromosomes 6, 8, 17, and 32, none of which had been previously reported. These regions emerged as promising candidates for future investigation through larger sample sizes and complementary approaches to confirm their potential role in VHSV resistance.

Interestingly, the QTL identified in our previous studies on chromosome 3 was not validated in the synthetic line by the present study. This QTL associated with resistance to VHSV in rainbow trout had been identified in two doubled haploid (DH) families (10), which had been derived from F0 breeders produced via mitogynogenetic reproduction and selected based on divergent viral replication in excised fin tissue (VREFT). Individuals exhibiting contrasting resistance phenotypes had been pair-mated to generate F1 isogenic crosses, each carrying one allele from each parent. Based on offspring survival and the contrast in VREFT values between breeders, two F1 crosses had been retained for QTL analysis.

The resulting F2 DH families combined the genomes of the two grandparents and were homozygous at all loci. This QTL detection relied on approximately one hundred polymorphic microsatellite markers, resulting in limited resolution for precise localisation. Within this constrained genetic background, the QTL on chromosome 3 accounted for 44% to 65% of the phenotypic variance in time to death following waterborne infection with VHSV strain 07.71, depending on the DH family. Additionally, the QTL explained 33% to 49% of the phenotypic variance in VREFT, supporting a key role for innate immune mechanisms in determining survival following VHSV infection. This QTL was subsequently validated in two additional DH families and a backcross between highly resistant and highly susceptible isogenic lines, and localised to the telomeric region of chromosome 3 (approximately the last 20 Mb) (11). This validation study used only eight markers within the QTL region, resulting in a limited resolution of its localisation. Nevertheless, collectively, it underscored the importance of the chromosome 3 QTL in resistance to waterborne VHSV infection.

As these studies had been conducted in highly constrained genetic backgrounds, either DH families or backcrosses between isogenic lines, their conclusions could not be generalised to more genetically diverse populations, such as wild or commercial stocks.

The synthetic line used in the present study encompasses a broader genetic background. It also represents the source population from which the previously developed doubled haploid (DH) families and isogenic lines were derived. As such, it provides an opportunity to assess the effects of the genomic regions identified in those earlier studies at the population level.

In the present study, no genomic region on chromosome 3 was detected as associated with progeny survival after balneation challenge in the synthetic population. While the VREFT is a good predictor of survival to balneation challenge (32), our data indicate that the resistance to VHSV does not rely solely on the mechanisms associated to genes on chromosome 3 which can impact viral replication, and point to additional key regions and mechanisms in the synthetic line. This is consistent with our observations across resistant isogenic lines, in which the mechanisms underlying VHSV resistance differed qualitatively and quantitatively between trout lines, supporting the polygenic basis of VHSV resistance (33,34).

Among the new suggestive regions associated with VHSV resistance some genes of interest were detected. Delta-like canonical Notch ligand 4 (DLL4, ENSOMYG00000064805 on chromosome 8), a transmembrane component of the Notch signalling pathway, is known to regulate a wide range of cellular processes in humans, including differentiation, proliferation, survival, angiogenesis, and vascular development. In addition to these roles, recent studies suggest that DLL4 contributes to immune regulation, particularly through modulation of macrophage polarization (35). Its expression is notably upregulated in activated human macrophages, where it promotes pro-inflammatory responses (36), supporting a potential role in antiviral defence mechanisms.

Zinc finger FYVE-type containing 1 (ZFYVE1, ENSOMYG00000003004 on chromosome 8) has been identified as a negative regulator of innate antiviral immunity. In black carp, it suppresses IRF3/IRF7-mediated antiviral responses (37), while in mammals it selectively inhibits MDA5-dependent signalling pathways, without affecting RIG-I signalling (38). Furthermore, ZFYVE1 has been shown to influence TLR3 signalling by facilitating ligand binding, thereby modulating downstream innate immune activation (39). These findings suggest a complex regulatory role in antiviral responses that could be relevant in the context of VHSV susceptibility.

NHS-like 3 (NHSL3, ENSOMYG00000053198 on chromosome 32) has also been implicated in inflammatory regulation, notably through its ability to potentiate TNFα–NFκB signalling, leading to enhanced pro-inflammatory responses (40). Such activity may contribute to shaping host immune responses during viral infection.

Mucin 4 (MUC4, ENSOMYG00000059070 on chromosome 6), a large glycoprotein and key structural component of mucus, plays a crucial role in forming protective epithelial barriers at interfaces with the external environment. By contributing to the physical barrier against pathogens, MUC4 may represent an important first line of defence, potentially limiting viral entry and infection.

Finally, in the suggestive regions, two members of the LDLR family are found to be genetically linked to VHSV survival, either *lrp3* (ENSOMYG00000058215 on chromosome 6) with intronic SNP in linkage disequilibrium with the top SNP on chromosome 6 or *lrp1* with intronic SNP as top SNP on chromosome 17.

While the functions of *lrp3* remains poorly known, it has been shown that *lrp3^-/-^*mice exhibit lower leukocyte count (41).

*lrp1* was selected as candidate gene for *in vitro* functional investigation given its role as entry receptor for a wide range of viruses (RVFV, SFTSV, DENV, OROV, YFV and JCV), including a rhabdovirus, the Chandipura virus (CHPV) (24,42–46). This highlights the role of *lrp1* as a multifunctional host factor that can facilitate viral attachment and internalisation across diverse viral families. Lastly, in mammals, *lrp1* was also involved in the regulation of immune responses, as it is associated with the inhibition of (TLR)-induced inflammation in macrophages (47).

However, our data showed that the *lrp1* paralog located on chromosome 2, in a region linked to VHSV resistance, is not an essential component of VHSV entry in CHSE cells, as viral RNA levels 30h post-infection were comparable between edited *lrp1^-/-^* clonal lines and wild-type (WT) cells. Indeed, several alternatives entry routes may exist for VHSV. In particular, tetraspanin CD9 has been demonstrated recently to act as an entry receptor in *Lateolabrax japonicus,* both *in vitro* and *in vivo* (48).

Although *lrp1* disruption did not affect viral RNA levels, phenotypic differences were observed in viral infectivity. One clonal line (C19) exhibited increased susceptibility compared to WT, with a 3.5-fold decrease in TCID_50_, whereas other edited lines showed no significant differences. Notably, in addition to knock-out of chromosome 2 and 22 *lrp1*, C19 carries heterozygous genotypes with mutations in two *lrp1* paralogs (WT/G367del on chromosome 7 *lrp1* and WT/G361_I362del on chromosome 16 *lrp1*). These two heterozygous mutations are located within the LDL receptor class B repeat (LDLR repeat B, IPR000033) of LRP1, one of the conserved cysteine-rich ligand-binding domains essential for ligand release and recycling of the receptor (49). This genetic configuration may reduce the level of *lrp1* gene product, resulting in haploinsufficiency, or produce a deleterious LRP1 protein with dominant-negative effects. Either mechanism could impair *lrp1* function and affecrt cellular homeostasis under VHSV-induced stress.

Moreover, the role of *lrp1* in modulating pro-inflammatory responses following VHSV infection was examined in the same mutant cell lines. The C19 clone, which displayed the most pronounced phenotype, also showed increased induction of pro-inflammatory markers. This observation is consistent with the known function of *lrp1* as a negative regulator of inflammatory signalling pathways. In contrast, the absence of phenotypic differences in clones C15 and C24 may be explained by compensatory effects from *lrp1* paralogs located on chromosomes 7 and 16 respectively, which remain unaltered (WT) in these clones. Further work will be required to clarify the functional interactions and immune functions between members of the *lrp1* family. Taken together, the results of the present study suggest that the association of SNPs within *lrp1* identified by GWAS is unlikely to reflect a direct role of this gene in epithelial cell susceptibility to VHSV entry. Instead, the data suggest a potential indirect role, whereby variation in *lrp1* may influence the fine regulation of pro-inflammatory responses, or other effects at the whole-organism level, contributing to differences in survival following VHSV infection in rainbow trout.

## Conclusion

The major QTL for VHSV resistance previously detected on chromosome 3 from experiments with DH lines was not confirmed when studying progeny survival after balneation challenge in the diverse synthetic rainbow trout population. Altogether, these data highlight the necessity of evaluating different phenotypes of resistance to virus. Integrating these findings with future GWAS results from other rainbow trout populations will help to understand the global genetic architecture of VHSV resistance.

Our study identified four novel suggestive regions using whole genome sequencing data, pointing to the gene *lrp1*, and other factors possibly affecting host virus interactions. While our in vitro *lpr1* mutants did not reveal obvious effect of resistance to VHSV, further experiments will be required to clarify the contribution of the genes located in the QTL identified here.

## Acknowledgements

This work was supported by INRAE, the French Ministry of Higher Education, Research and Innovation and AAP Carnot France Futur Elevage 2023 (RESSHV project). We thank the technical staff of IERP for their help with the rainbow trout waterborne challenge to VHSV (IERP-UE907 DOI: 10.15454/1.5572427140471238E12, INRAE Jouy-en-Josas, France).

## Supporting information

**S1 Table. List of the 13 rainbow trout-specific microsatellites used for genotyping**

**S2 Table. Pairwise Probability of Identity by State among the 14 whole genome sequenced individuals**

**S3 Table. qPCR primers used in this study**

**S4 Table. SNP in strong linkage disequilibrium (r²** ≥ **0.85) within a 1 Mb window (±500 kb around every suggestive SNP)**

**S5 Table. Sequences of each CHSE-EC-*lrp1-/-* cell line at the *lrp1* loci**

